## Supplementary Material for "Convergence of resistance and evolutionary responses in *Escherichia coli* and *Salmonella enterica* co-inhabiting chicken farms in China"

Michelle Baker *et al*

**This PDF file includes:**

Figs. S1 to S13

**Other Supplementary Materials for this manuscript include the following:**

Tables S1 to S7

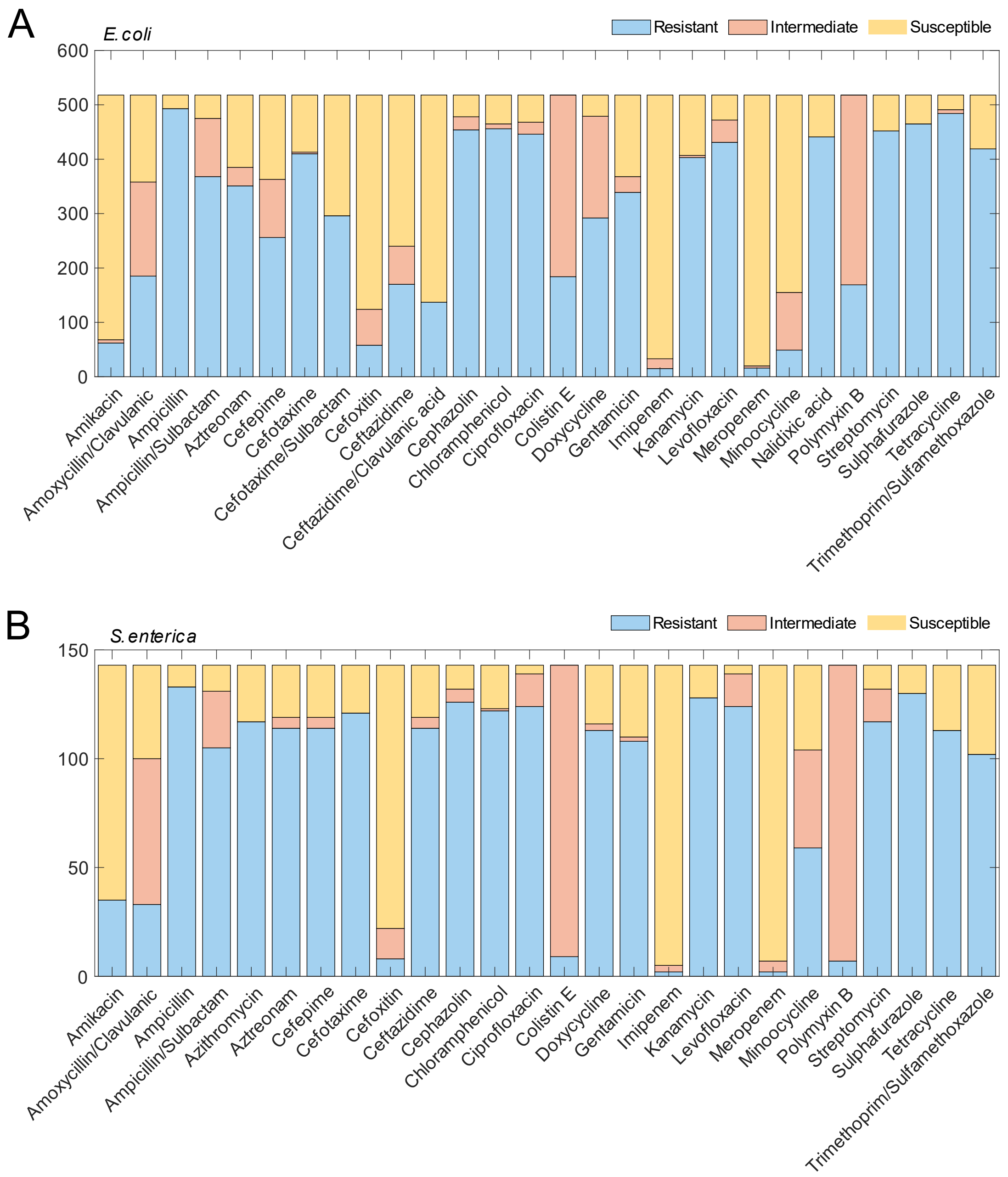


**Fig. S1**. Number of susceptible, intermediate, and resistant *E. coli* and *S. enterica* isolates, based on antimicrobial susceptibility testing by broth microdilution. (A) AMR phenotypes for 518 *E. coli* isolates tested against a panel of 28 antibiotics. (B) AMR phenotypes for 143 *S. enterica* isolates tested against a panel of 26 antibiotics.

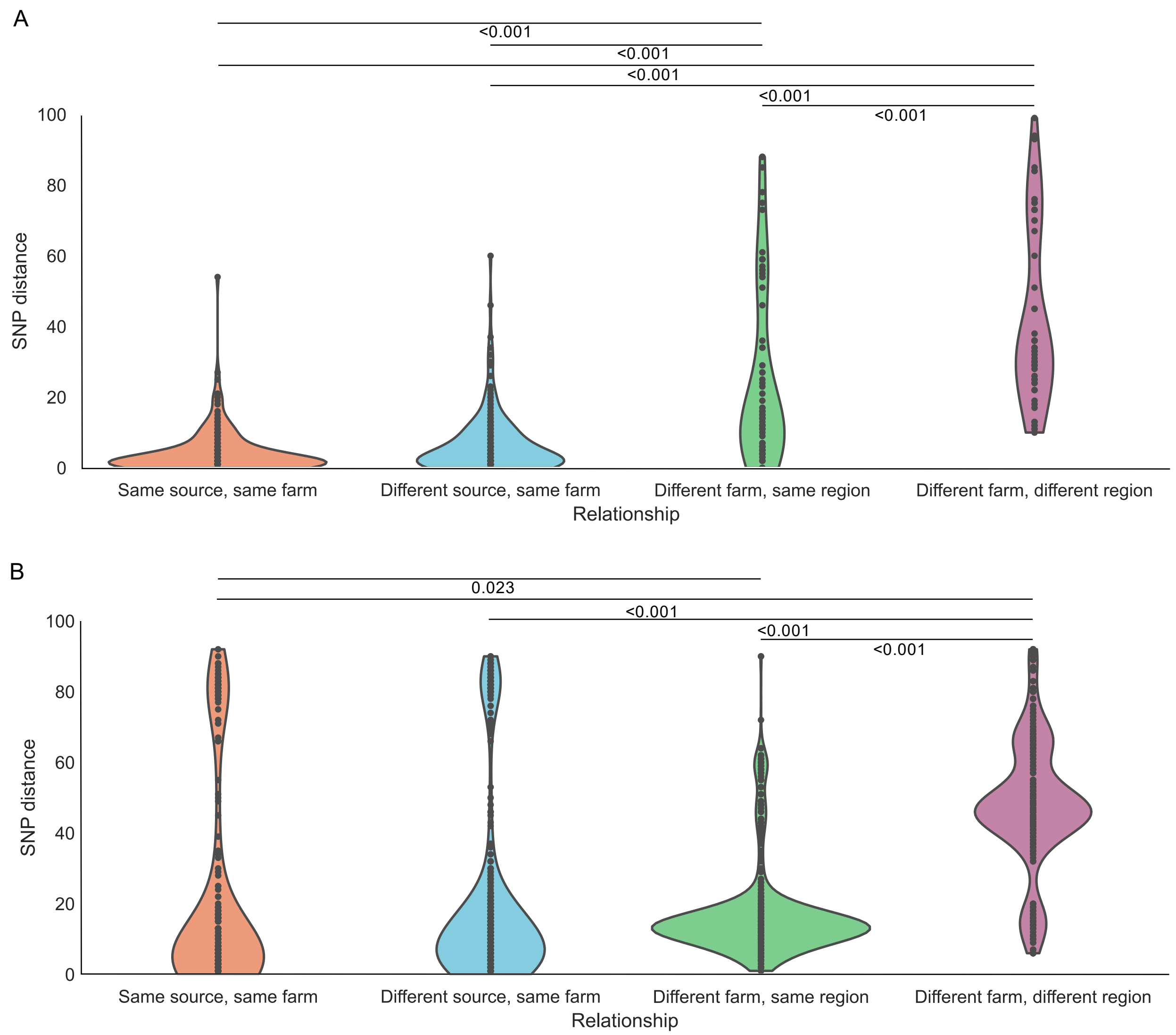


**Fig. S2**. Distribution of pairwise comparisons of SNP distance between (A) *E. coli* and (B) *S. enterica* isolates across source types, different farms, and provinces. Only pairs with less than 100 SNPs were included in the analysis *(19)*.

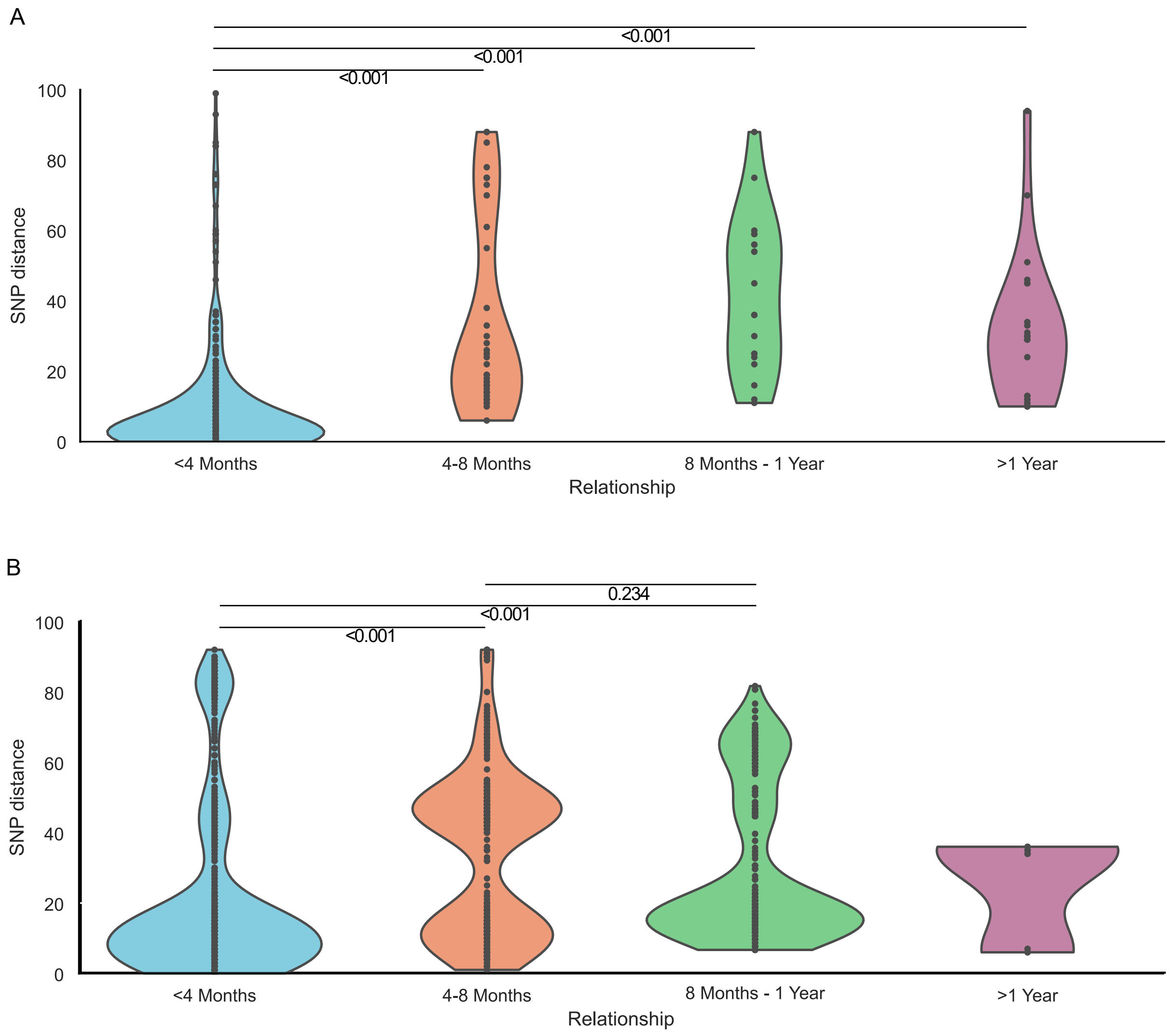


**Fig. S3**. Distribution of pairwise comparisons of SNP distance between (A) *E. coli* and (B) *S. enterica* isolates across collection dates. Only pairs with less than 100 SNPs were included in the analysis *(19)*. The distribution categories refer to the difference in dates between each isolate pair.

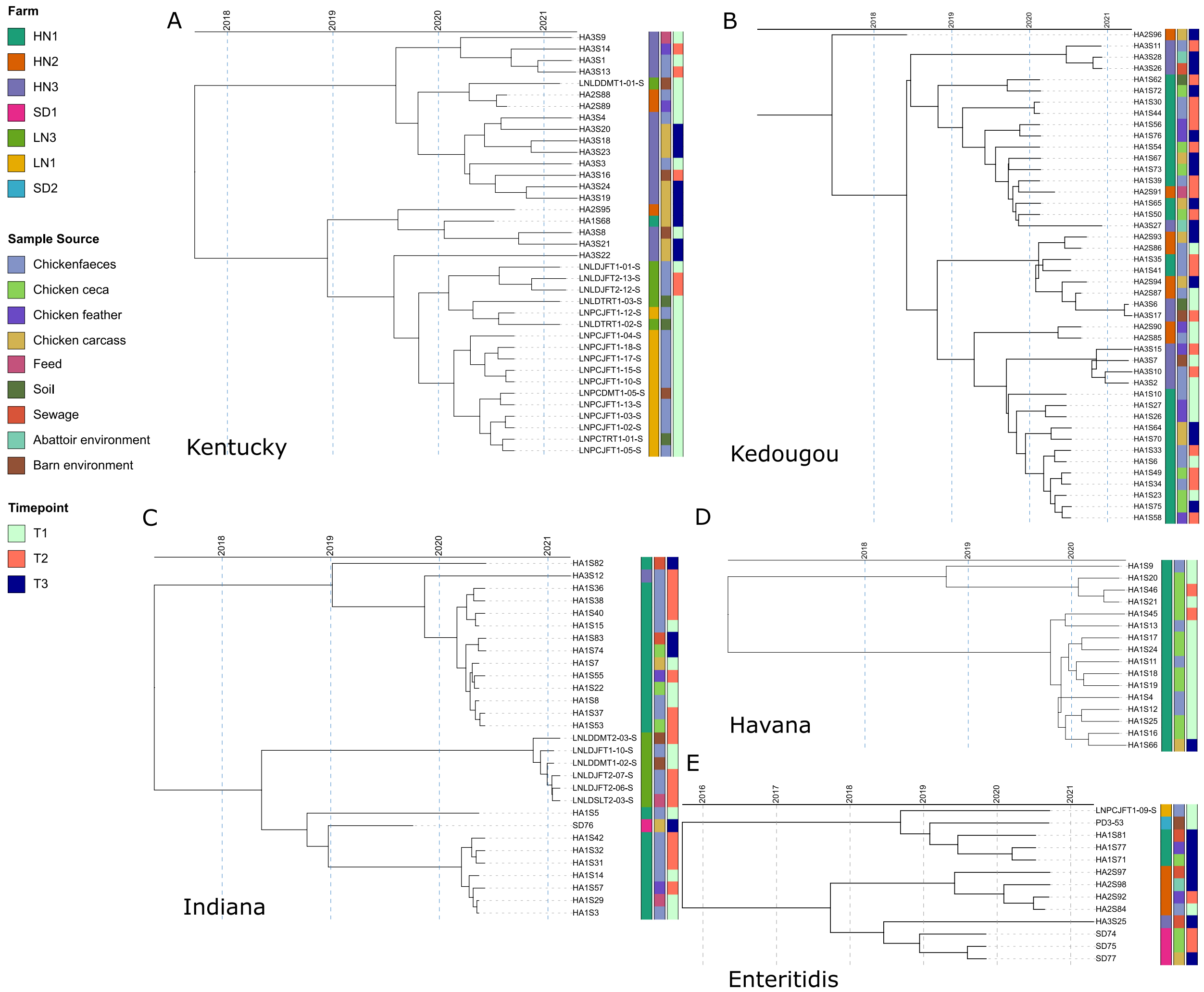


**Fig S4.** Bayesian divergence analysis of *S. enterica* isolates in five serotypes (A) Kentucky; (B) Kedougou; (C) Indiana; (D) Havana and (E) Enteritidis. The sample source and the farm the isolates were taken from and the timepoint for each sample are displayed as coloured strips.

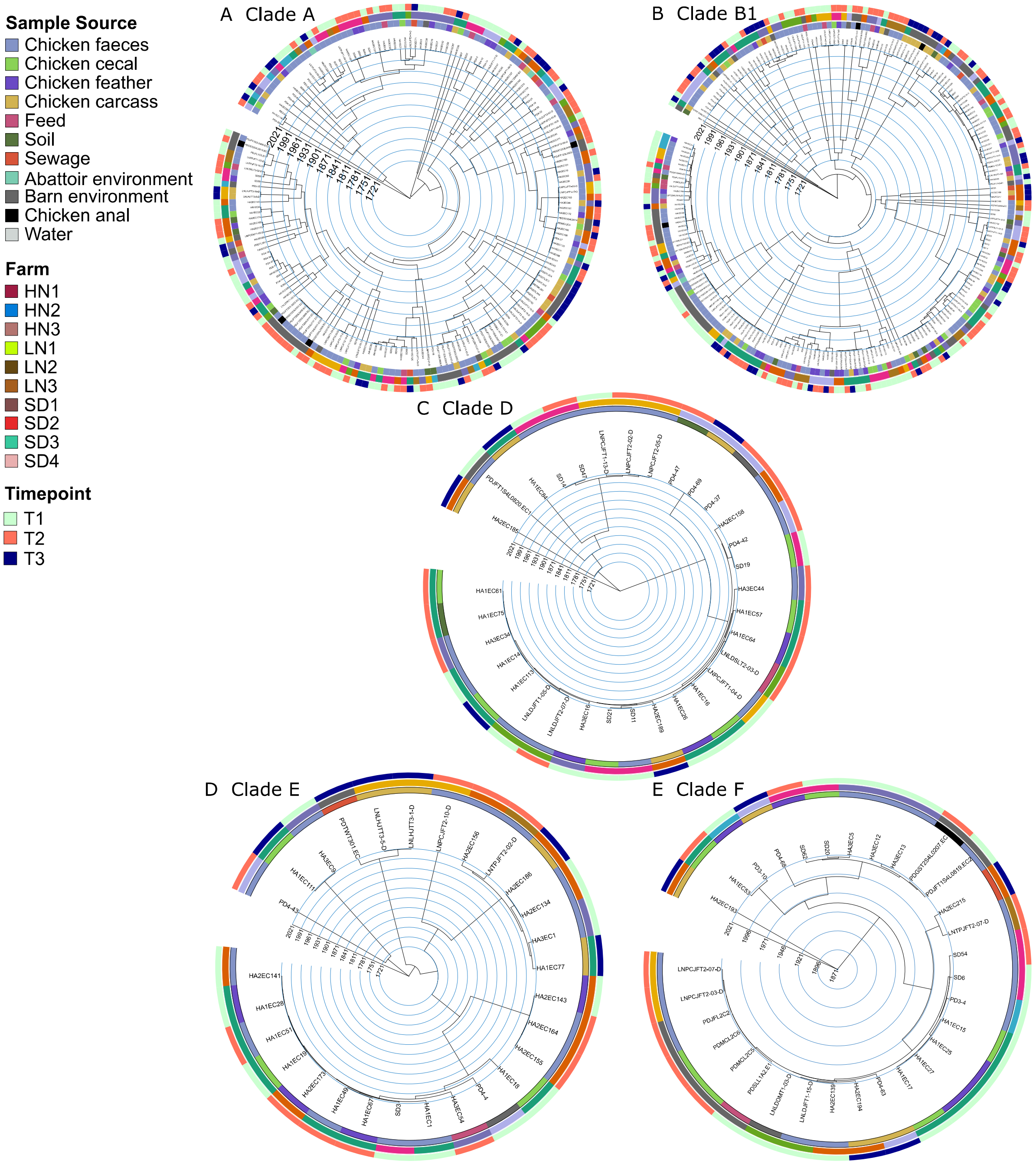


**Fig S5.** Bayesian divergence analysis of *E. coli* isolates in five phylogroups (A) Clade A; (B) Clade B1; (C) Clade D; (D) Clade E and (E) Clade F. Sample source and the farm the isolates were taken from and the timepoint for each sample are displayed as coloured rings.

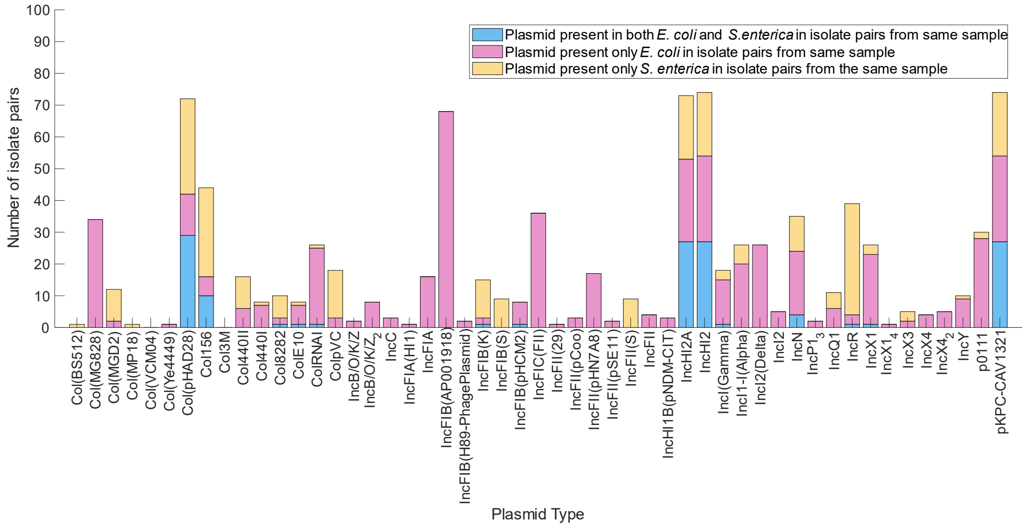


**Fig S6.** Plasmid presence in *E. coli* and *S. enterica* isolate pairs from the same samples. Number of isolate pairs with the same plasmids in both *E. coli* and *S. enterica* (blue), *E. coli* only (pink) and *S. enterica* only (yellow), separated by replicon type. Only isolates where both species were isolated from the same sample were included in this analysis (i.e. 113 *E. coli* and 113 *S. enterica* isolates).

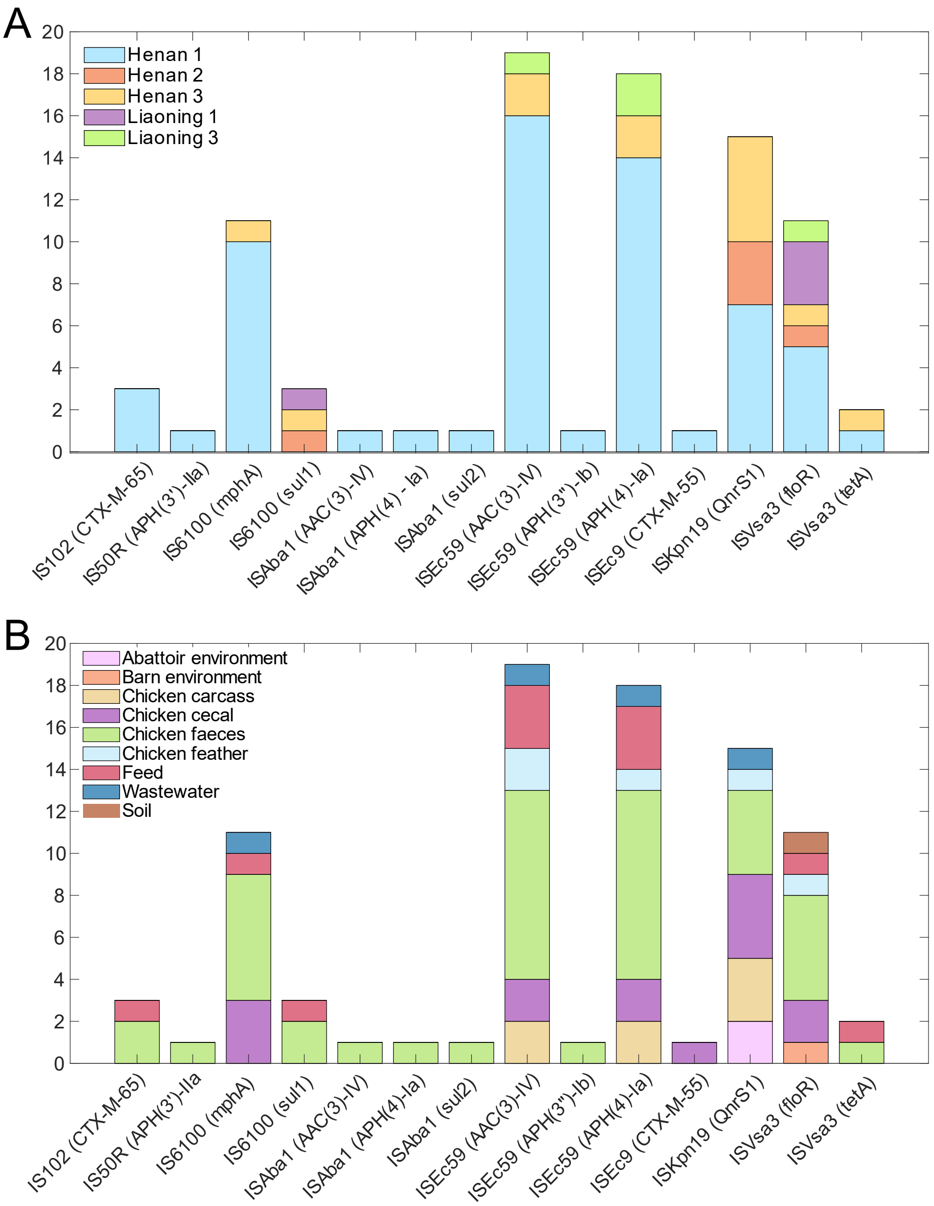


**Fig S7.** Numbers of different mobile ARGs found in isolate pairs of *S. enterica* and *E. coli* taken from the same samples. Of 113 *E. coli* and *S. enterica* isolates pairs collected from the same samples, 88 were found to have the same mobile ARG. (A) Stacked bar plot showing the breakdown, by farm, of mobile ARGs found in both *E. coli* and *S. enterica* isolates. (B) Stacked bar plot showing the breakdown by source type of mobile ARGs found in both *E. coli* and *S. enterica* isolates.

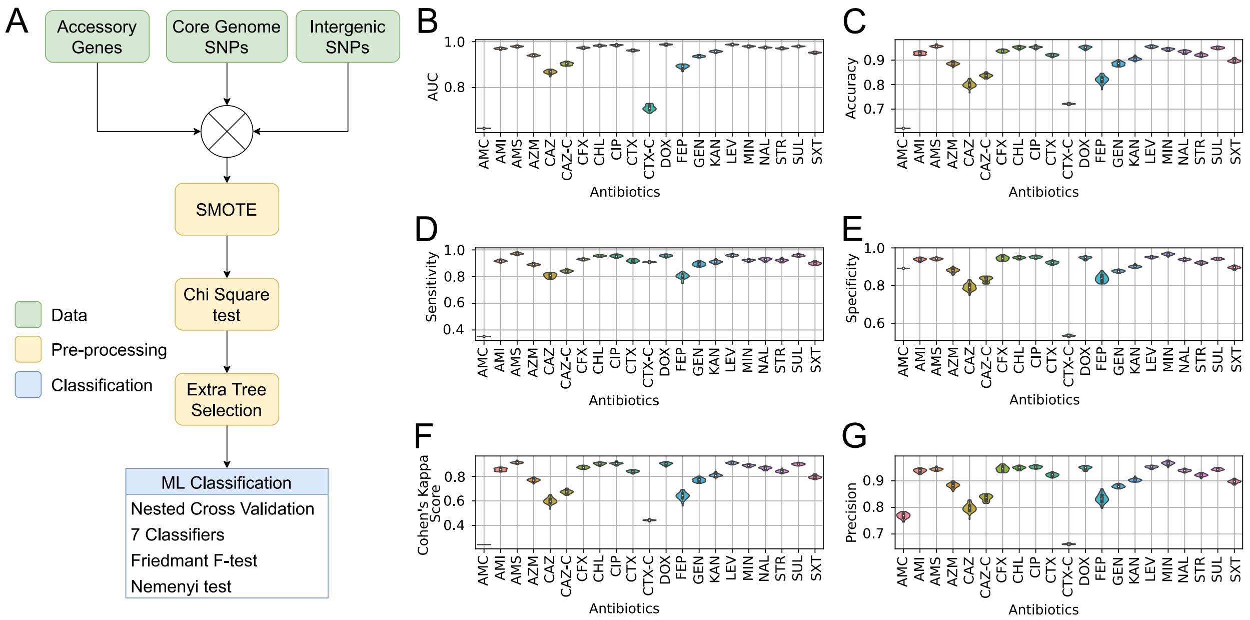


**Fig S8.** Supervised machine learning pipeline accurately predicts *E. coli* resistance susceptibility profiles. (A) Flow diagram showing machine learning pipeline including data (green), pre-processing steps (yellow) and classification (blue). (B) – (G) Machine learning performance results for six performance indicators: (B) area under the curve AUC, (C) accuracy, (D) sensitivity, (E) Specificity, (F) Cohen's kappa score, and (G) precision from 30 training runs for each antimicrobial. The results shown are for the best classifier Random Forest, as defined by the Nemenyi test (Fig. S10A). Predictive models were generated for twenty-one different antimicrobials (X axis): amoxycillin/clavulanic acid (AMC), amikacin (AMI), ampicillin/sulbactam (AMS), aztreonam (AZM), ceftazidime (CAZ), ceftazidime/clavulanic acid (CAZ-C), cefoxitin (CFX), chloramphenicol (CHL), ciprofloxacin (CIP), cefotaxime (CTX), cefotaxime/ clavulanic acid (CTX-C), doxycycline (DOX), cefepime (FEP), gentamicin (GEN), kanamycin (KAN), levofloxacin (LEV), minocycline (MIN), nalidixic acid (NAL), streptomycin (STR), sulfisoxazole (SUL), and trimethoprim/sulfamethoxazole (SXT).

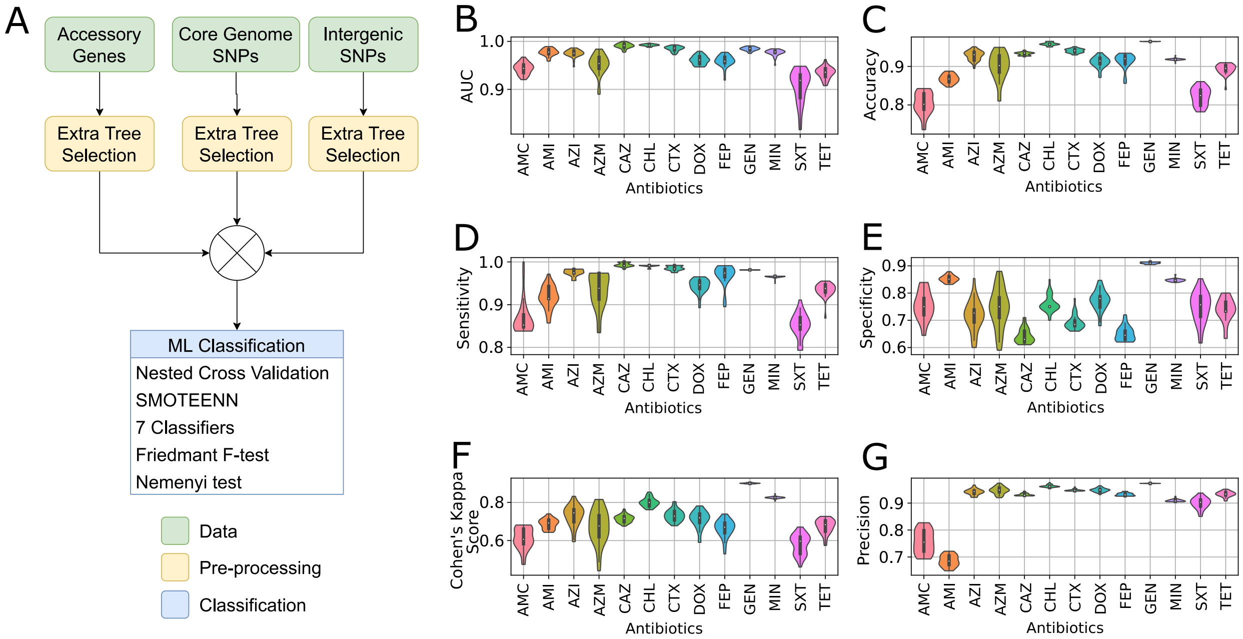


**Figure S9.** Supervised machine learning pipeline accurately predicts *S. enterica* resistance susceptibility profiles. (A) Flow diagram showing machine learning pipeline including data (green), pre-processing steps (yellow) and classification (blue). (B) – (G) Machine learning performance results for six performance indicators: (B) area under the curve AUC, (C) accuracy, (D) sensitivity, (E) Specificity, (F) Cohen's kappa score, and (G) precision from 30 training runs for each antimicrobial. The results shown are for the best classifier Linear SVM, as defined by the Nemenyi test (Fig. S10B). Predictive models were generated for thirteen different antimicrobials (X axis): amoxycillin/clavulanic acid (AMC), amikacin (AMI), azithromycin (AZI), aztreonam (AZM), ceftazidime (CAZ), chloramphenicol (CHL), cefotaxime (CTX), doxycycline (DOX), cefepime (FEP), gentamicin (GEN), minocycline (MIN), trimethoprim/sulfamethoxazole (SXT), and tetracycline (TET).

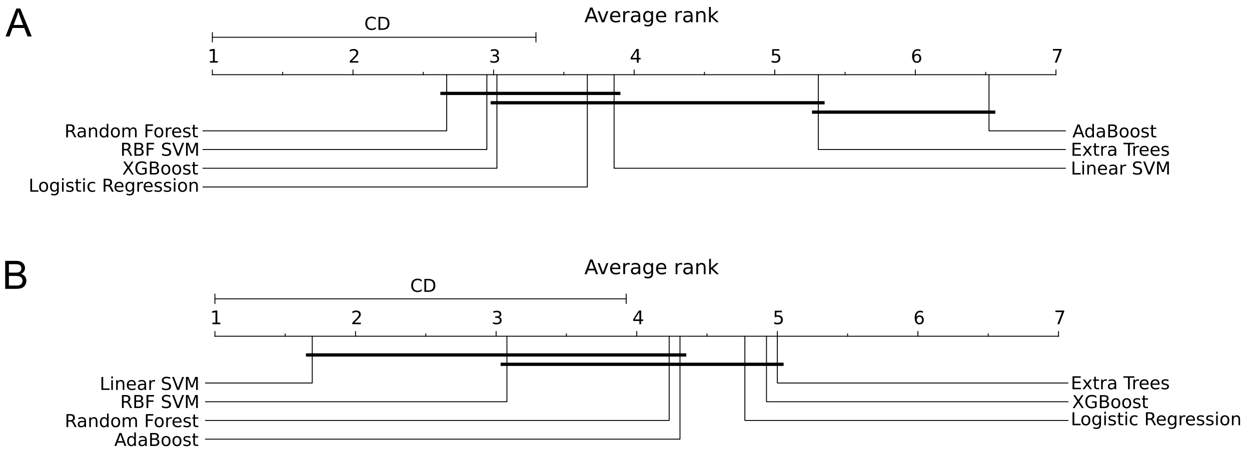


**Fig S10.** Nemenyi *post-hoc* tests *(89)*. Comparison of the performance of the 5 classifiers and 2 meta-methods, using their average ordinal rank over the 11 antibiotics analysed based on the AUC performance metric for (A) *E. coli* and (B) *S. enterica*. The x-axis indicates the average ordinal rank of the machine learning methods. The scale is from 1 (best rank) to 7 (worst rank). The ordinal rank of a classifier is defined as follows: the ML method with the best AUC is given rank 1, the second-best AUC rank 2 and the *n*-th AUC best rank *n*, with *n* being the number of machine learning methods used. For each antibiotic, the methods are ranked between 1 (highest AUC) and 7 (lowest AUC), since in this case there are 7 machine learning methods used. Next, for each method, the ranks are averaged based on the 11 antibiotics studied. The critical distance (CD) is defined based on the Nemenyi *post-hoc* test, all the methods that fall in the same bold bar below the axis are considered statistically equivalent based on the CD value *(89)*.

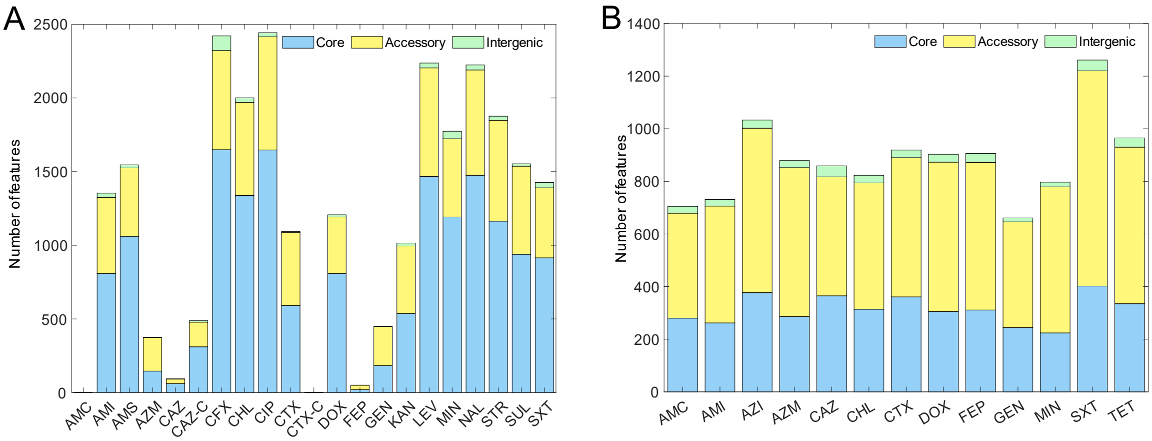


**Fig S11.** Number of features correlated to resistance-susceptibility profiles by ML. For each antibiotic model for (A) *E. coli* and (B) *S. enterica*, the number of core (blue), accessory (yellow) and intergenic (green) features are shown. The antibiotics shown are amoxicillin/clavulanic acid (AMC), amikacin (AMI), ampicillin/sulbactam (AMS), azithromycin (AZI), aztreonam (AZM), ceftazidime (CAZ), ceftazidime/clavulanic acid (CAZ-C), cefoxitin (CFX), chloramphenicol (CHL), ciprofloxacin (CIP), cefotaxime (CTX), cefotaxime/ clavulanic acid (CTX-C), doxycycline (DOX), cefepime (FEP), gentamicin (GEN), kanamycin (KAN), levofloxacin (LEV), minocycline (MIN), nalidixic acid (NAL), streptomycin (STR), sulfisoxazole (SUL), trimethoprim/sulfamethoxazole (SXT), and tetracycline (TET).

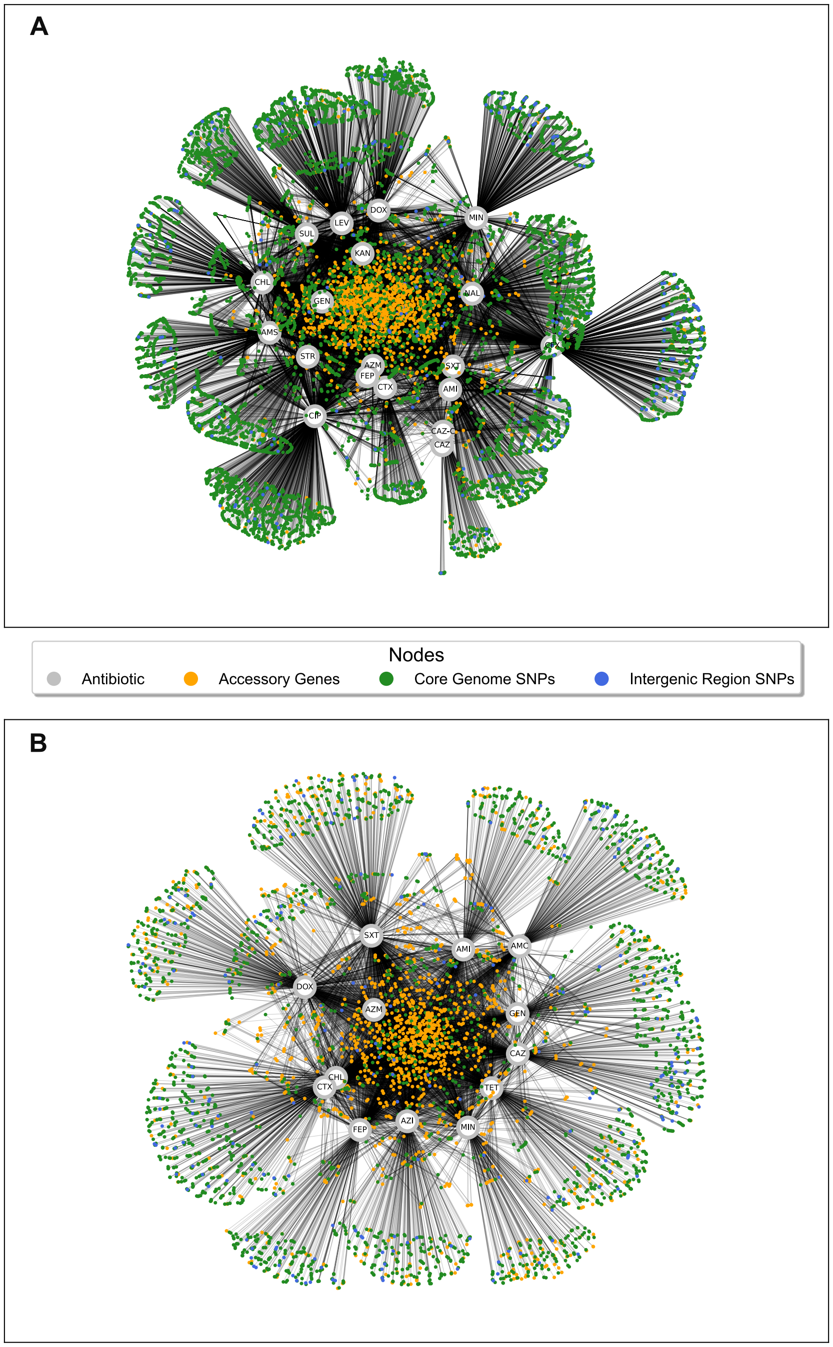


**Fig S12.** Undirected graph network indicating the genomic features found for each antibiotic model for (A) *E. coli* and (B) *S. enterica*. The colour of the node indicates which genomic feature it belongs to accessory genes are in orange, core genome SNPs in green and intergenic region SNPs in blue. The antibiotic models are coloured in grey, and indicated as three letter antibiotic abbreviations.

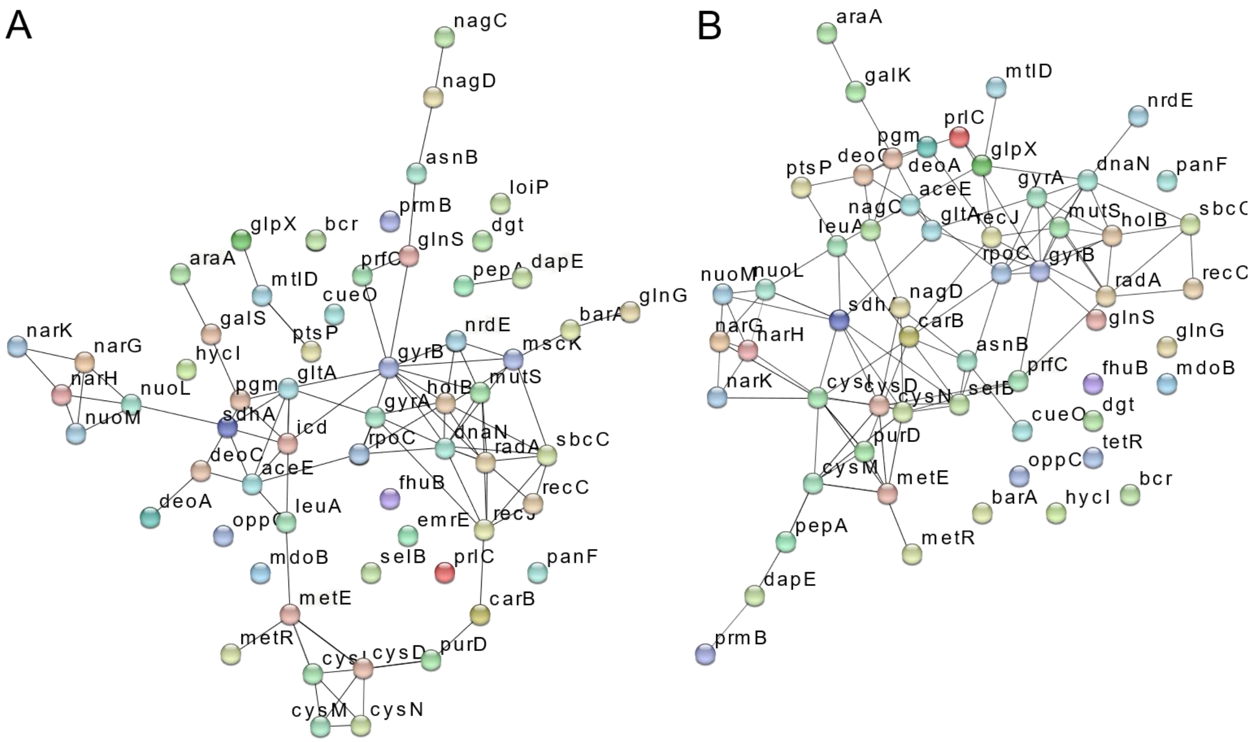


**Figure S13.** The most significant genes correlated to AMR profiles, as predicted by ML, in both species show high levels of protein-protein interaction. Genes that were in the top 10% of features (ranked according to the maximum feature importance in 30 runs) in both species, *E. coli* and *S. enterica* are plotted in a protein-protein interaction network, according String database interactions (confidence threshold 0.4). (A) genes plotted according to interactions in *E. coli K12 MG1655*. (B) genes plotted according to interactions in *S. enterica subsp. enterica serovar Typhimurium str. LT2*. Network generated by StringDB v11.5.

**Table S1 (Separate File)**

Characteristics of the *E. coli* and *S. enterica* isolates, including collection details, typing data, assembly statistics, genomic features, and results of antibiotic susceptibility testing and CARD ARGs.

**Table S2. (Separate File)**

Presence-absence of plasmids, separated by replicon type, on each of the 518 *E. coli* and 143 *S. enterica* isolates by comparison to the PlasmidFinder database with 80% identity and 70% coverage thresholds.

**Table S3. (Separate File)**

List of the shared mobile ARGs within the 113 isolate pairs of *E. coli* and *S. enterica* isolates cultured from the same sample. Mobile ARGs are defined as the presence of and MGE within 5kb upstream or downstream of the known ARG. Isolate ID, mobile ARG, farm and source type are given.

**Table S4. (Separate File)**

Machine learning performance for each of the 7 classification methods used for both *E. coli* isolates and *S. enterica* isolates correlating the genetic features (SNPs and accessory genes) to the resistance/susceptibility against a panel of antimicrobials. The performance metrics were accuracy (TP+TN/(P+N)), sensitivity (true positive rate: TP/P), specificity (true negative rate: TN/N), AUC and precision. The scores for each performance metric were computed from 30 simulations using nested cross-validation. The mean ± standard deviation of these 30 iterations was then used as the result statistics for the performance.

**Table S5. (Separate File)**

Feature importance of genes selected by the ML pipeline as correlated to the resistance-susceptibility of either *E. coli* or *S. enterica* to a panel of 26 antibiotics. The genes listed were either accessory genes or core genes containing a SNP correlated to either resistance or susceptibility. Where the gene was identified as a known AMR gene present in public databases (by comparison using BLAST) the accession is given. Feature importance is given as average importance across models. For each model, the feature importance was calculated based on the Gini Importance of an ExtraTree Classifier with 50 estimators.

**Table S6. (Separate File)**

KEGG ontology of the 88 genes found in the top 10% of most important features in both the *E. coli* and *S. enterica* models.

**Table S7. (Separate File)**

Genes used as input to the genome scale models, derived from the top 10% of features in each of the machine learning models.
